# SingleRust: A High-Performance Toolkit for Single-Cell Data Analysis at Scale

**DOI:** 10.1101/2025.08.04.668429

**Authors:** Ian F. Diks, Matthias Flotho, Andreas Keller

## Abstract

Single-cell RNA sequencing studies increasingly generate datasets exceeding 10 million cells, surpassing the memory capacity of standard analytical tools on typical institutional infrastructure. Here we introduce SingleRust, a computational framework that addresses these constraints through systematic algorithmic optimizations and systems-level design. Key improvements include sparse masked principal component analysis that reduces memory footprint while preserving biological signal, lock-free parallel implementations for differential expression testing, and adaptive k-nearest neighbor algorithms that automatically select optimal data structures based on dataset size. These optimizations achieve 2.4-25.5-fold performance improvements and 1.3-3.0-fold memory reduction compared to Scanpy, enabling routine analysis of 30 million cells on our representative test system with 512 GB RAM. Comprehensive validation confirms numerical equivalence with established methods while maintaining biological interpretation fidelity. SingleRust maintains full compatibility with the AnnData ecosystem while providing researchers immediate access to population-scale analyses on existing infrastructure, addressing a critical bottleneck in single-cell genomics workflows.

## 1 Introduction

Single-cell RNA sequencing has become an essential technology for dissecting cellular heterogeneity in complex biological systems. Applications span from comprehensive cell atlas projects [1] to population-scale disease studies [2] and emerging clinical diagnostics [3]. As the technology matures, datasets have grown exponentially, from thousands of cells in early studies to over 100 million cells in current flagship projects [4, 5]. This growth trajectory shows no signs of slowing, with spatial transcriptomics and longitudinal clinical monitoring poised to generate even larger datasets [6].

Current computational infrastructure has not scaled proportionally with data generation capabilities. The field’s primary analytical frameworks, Seurat [7] and Scanpy [8], were architected when thousand-cell datasets were standard and million-cell datasets were exceptional. These tools now encounter fundamental limitations when processing contemporary datasets [9]. On typical institutional infrastructure with 512 GB RAM—representing accessible compute resources for most research groups—current tools face memory exhaustion, performance degradation, or require algorithmic compromises that may impact biological discovery [9].

The scalability challenge extends beyond simple memory constraints to fundamental architectural limitations inherent in interpreted languages. Python’s Global Interpreter Lock (GIL) serializes execution, preventing effective utilization of multi-core processors that have become standard in modern computing infrastructure. Dynamic typing necessitates runtime type checking and reference counting for every numerical operation, introducing computational overhead that compounds with dataset size. Garbage collection creates non-deterministic performance characteristics, making resource planning difficult and potentially interrupting time-sensitive analyses. These architectural constraints become particularly acute for sparse matrix operations—the dominant data structure in single-cell analysis—where interpreted languages must manage millions of individual object allocations.

Emerging applications demand computational frameworks that provide predictable performance and efficient resource utilization. Population genetics studies require analyzing millions of cells across dozens of individuals to capture human diversity. Clinical applications necessitate deterministic execution for regulatory compliance, guaranteed completion times for time-sensitive decisions, and robust error handling to prevent silent failures. Current frameworks, designed for exploratory research, struggle to meet these requirements without substantial modifications that often compromise performance or functionality.

Systems programming languages offer architectural advantages that could address these limitations. Compiled languages eliminate interpretation overhead and enable compile-time optimizations specific to numerical workloads. Zero-copy semantics avoid redundant memory allocations common in interpreted languages. True parallelization without global locks enables efficient utilization of modern multi-core processors. Static typing and compile-time memory safety checks prevent entire classes of runtime errors. These characteristics suggest that reimplementing core single-cell algorithms in a systems programming language could overcome current scalability barriers.

We recognize that the single-cell analysis ecosystem has evolved sophisticated solutions for handling large datasets. Scanpy offers backed processing for out-of-memory datasets and Dask-based parallelization, while Rapids-Single-Cell provides GPU acceleration for compatible hardware. These tools excel at rapid prototyping, leveraging Python’s extensive ecosystem, and iterative exploration—capabilities that remain essential for biological discovery. Our work addresses a complementary niche: high-performance, high-throughput pipelines that prioritize in-memory processing with predictable performance, minimal resource consumption, and deterministic execution. This specialization reflects the reality that different analytical contexts demand different computational trade-offs.

Here we present SingleRust, a computational framework that applies systems programming principles to single-cell analysis. Built on Rust’s ownership model and zero-copy semantics, SingleRust reimplements six essential single-cell operations: quality control filtering, count normalization, highly variable gene identification, principal component analysis, differential expression testing, and k-nearest neighbor graph construction. The framework maintains full compatibility with the AnnData format and provides APIs consistent with existing tools, enabling incremental adoption within established workflows. Our benchmarks use standard configurations without GPU or distributed computing optimizations, reflecting typical institutional deployments where such resources may be limited or require complex implementation. We systematically evaluate whether compiled-language architectures can address the scalability and reliability requirements of next-generation single-cell studies while preserving the numerical fidelity required for biological discovery.

## 2 Results

We evaluated SingleRust’s performance against Scanpy using datasets ranging from 10,000 to 30 million cells on standard 512 GB RAM systems. Benchmarks encompassed six core single-cell operations under matched parameter conditions. SingleRust successfully processed 30 million cell datasets, three times the practical limit observed with current tools—while demonstrating runtime improvements of 2.4 to 25.5-fold and memory reductions of 1.3 to 3.0-fold across operations (Figure 2). These performance gains were achieved while maintaining numerical equivalence with established methods, as detailed in subsequent validation analyses.

**Fig. 1.**
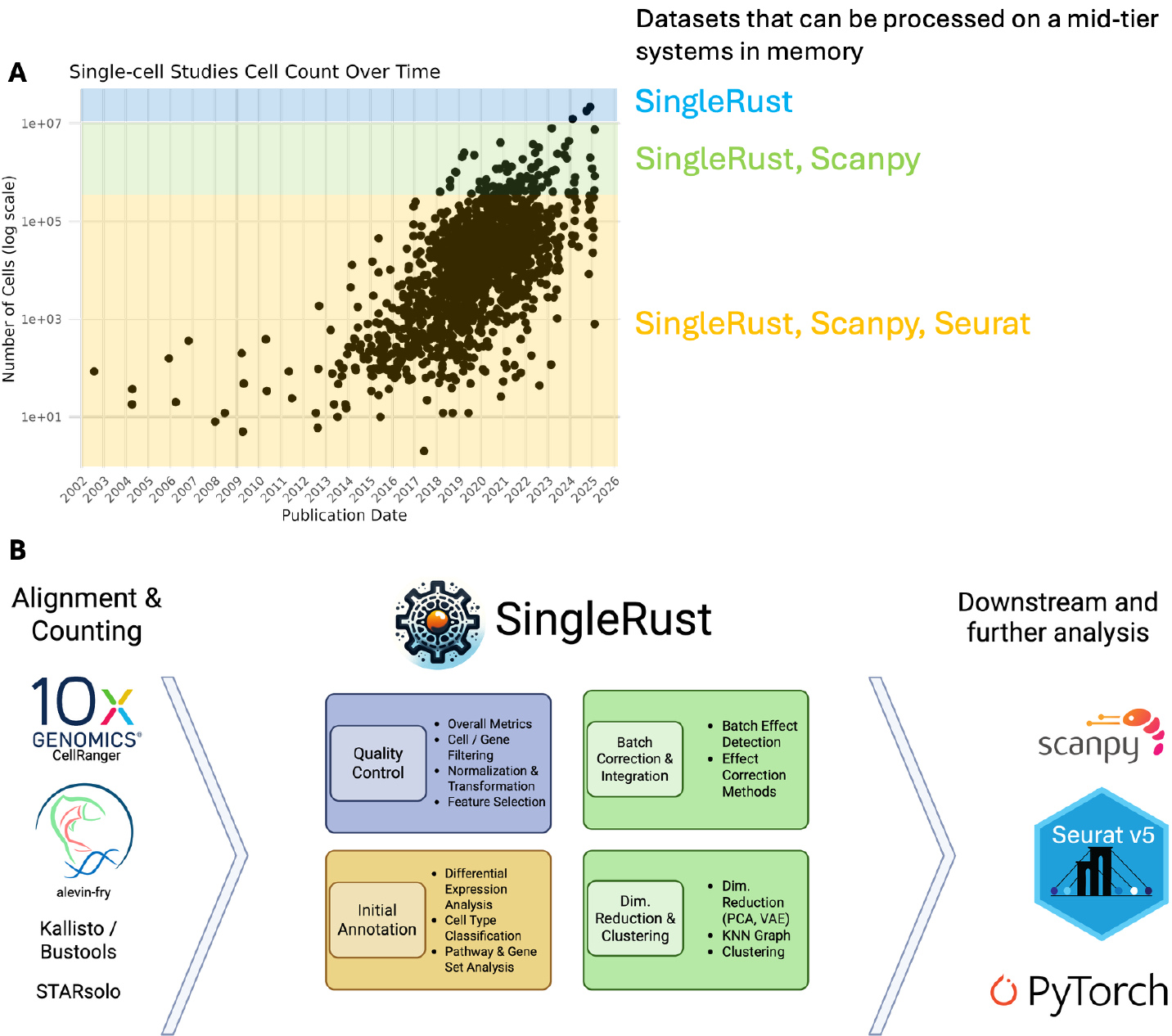
Exponential growth of single-cell datasets and the SingleRust solution. **(a)** Single-cell dataset sizes from 2002-2025 (log scale) with colored regions showing computational limits on 512 GB RAM: yellow (all tools, *<*1M cells), green (SingleRust/Scanpy only, 1-10M cells), blue (SingleRust only, 10-20M+ cells). **(b)** SingleRust workflow integrating with existing single-cell analysis pipelines through four core modules: Quality Control, Initial Annotation, Batch Correction & Integration, and Dimensionality Reduction & Clustering.

**Fig. 2.**
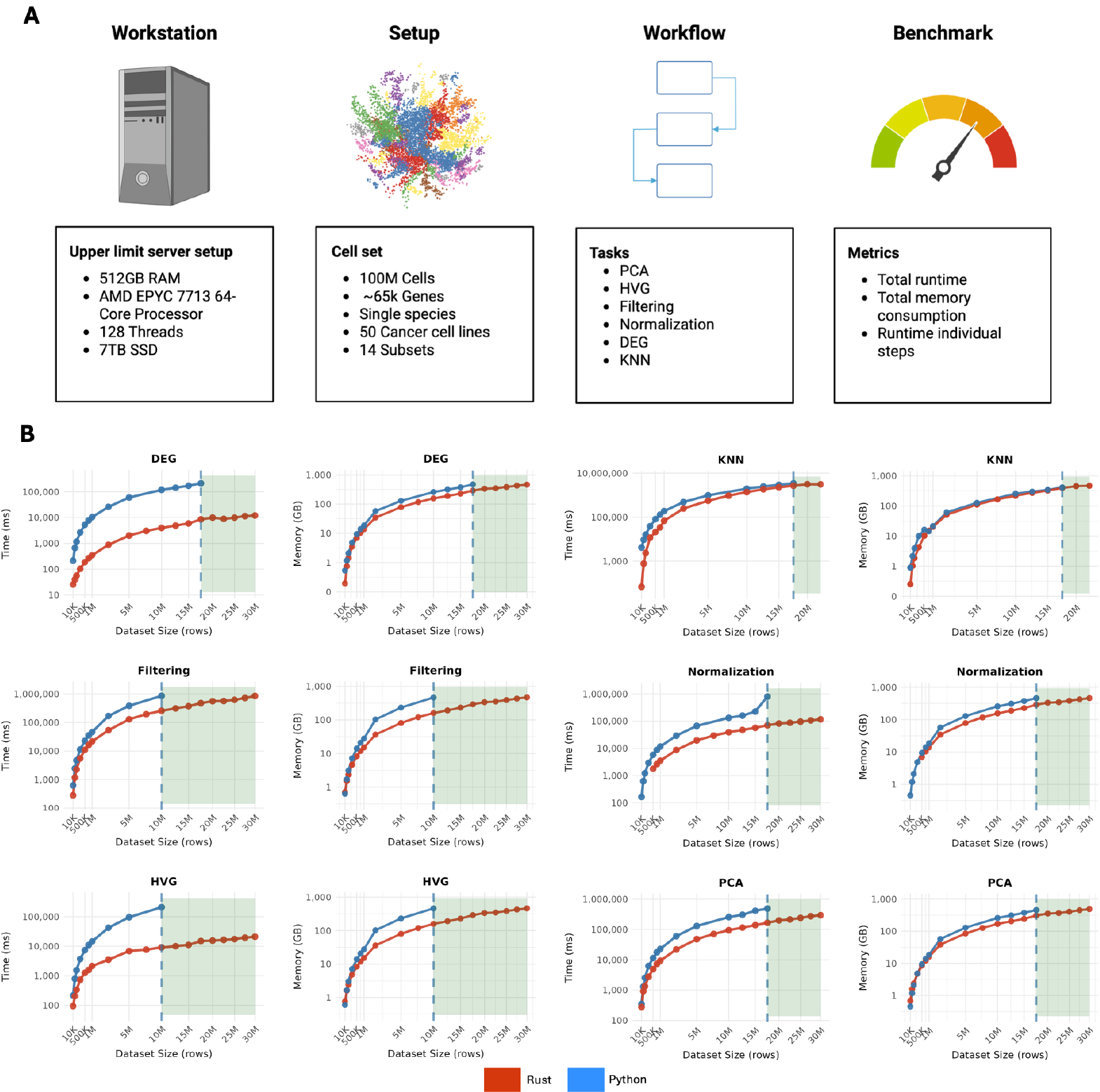
Rust Delivers Significant Performance Improvements over Python. **(A)** Benchmarking setup showing the computational environment (512GB RAM server), dataset characteristics (Tahoe 100M cells), analysis workflow (six core operations), and evaluation metrics. **(B)** Runtime and memory consumption comparison between SingleRust (blue) and Scanpy (red) across six essential single-cell operations, arranged in two columns with three operations each. For each operation, performance (runtime in milliseconds, log scale) and memory usage (GB, log scale) are shown side by side. X-axis shows dataset size from 10k to 20M cells. Green shaded area indicates dataset sizes typically inaccessible to Scanpy. Dashed line indicates the cutoff on dataset size for each operation.

### 2.1 SingleRust Achieves Order-of-Magnitude Performance Gains

SingleRust outperformed Scanpy by 2.2 to 24.6-fold across six core single-cell operations (Figure 2B). The largest gains emerged in computationally intensive tasks: differential expression analysis accelerated 25.5x, while k-nearest neighbor graph construction demonstrated substantial improvements of 7.9x on average, with particularly strong performance on smaller datasets (36x speedup for 10,000 cells). Highly variable gene identification ran 8.5x faster, and normalization showed 4.3x improvement. Even heavily optimized operations benefited—PCA accelerated 2.4x despite Scanpy’s existing optimizations, while memory-bandwidth-limited filtering improved 2.4x. Notably, cold-start performance showed even more dramatic improvements, with KNN achieving up to 60.9x speedup, highlighting SingleRust’s efficient memory initialization compared to Python’s object allocation overhead.

These gains stem from Rust’s fundamental advantages over interpreted languages. Zero-copy semantics eliminate Python’s object duplication overhead, enabling direct operation on sparse matrices. Lock-free parallelization—unconstrained by Python’s global interpreter lock—allows concurrent processing across CPU cores. Combined with algorithm-specific optimizations for sparse data structures, these architectural choices transform computational bottlenecks. Tasks that require hours in Python complete in minutes, making population-scale analyses practical on standard infrastructure.

### 2.2 Substantial Reduction in Resource Consumption

Beyond runtime improvements, SingleRust consistently reduces memory consumption compared to Scanpy, achieving an average 1.6-fold improvement across all operations (Figure 2C). While some operations showed substantial gains—up to 3.0-fold reduction for HVG identification and filtering, others like KNN and PCA demonstrated modest but meaningful improvements of approximately 30%. DEG and normalization operations showed 1.7x reduction. The memory savings result from several technical factors. Zero-copy semantics avoid data duplication during operations. The absence of Python’s global interpreter lock reduces synchronization overhead. Additionally, we implemented optimizations such as on-demand type conversion—when operations require higher precision, we convert individual values rather than entire matrices, reducing memory allocation. These consistent memory reductions across all operations enable processing of larger datasets on the same hardware. Notably, even at 30 million cells, SingleRust remained well within system capacity—HVG used 463 GB, PCA used 487 GB, and KNN peaked at 466 GB on our 512 GB system, leaving 40 GB available.

### 2.3 Numerical Accuracy and Statistical Equivalence with Established Methods

The performance and memory improvements achieved by SingleRust would be meaningless without maintaining numerical accuracy. To ensure our optimizations preserve scientific validity, we systematically validated all operations against Scanpy’s established implementations.

### HVG Validation

Despite the 8.5x speedup and 3.0x memory reduction, SingleRust produces identical results to Scanpy. Testing with a limit of 2,500 highly variable genes, both implementations identified exactly the same gene set with perfect overlap (Figure 3B). The underlying statistics, normalized dispersion and mean expression, showed perfect correlation (R = 1.0) with zero mean difference. This numerical equivalence persisted across multiple dataset sizes (100k, 500k, 1M cells) and remained consistent between runs, confirming that our memory-efficient algorithms preserve computational accuracy.

**Fig. 3.**
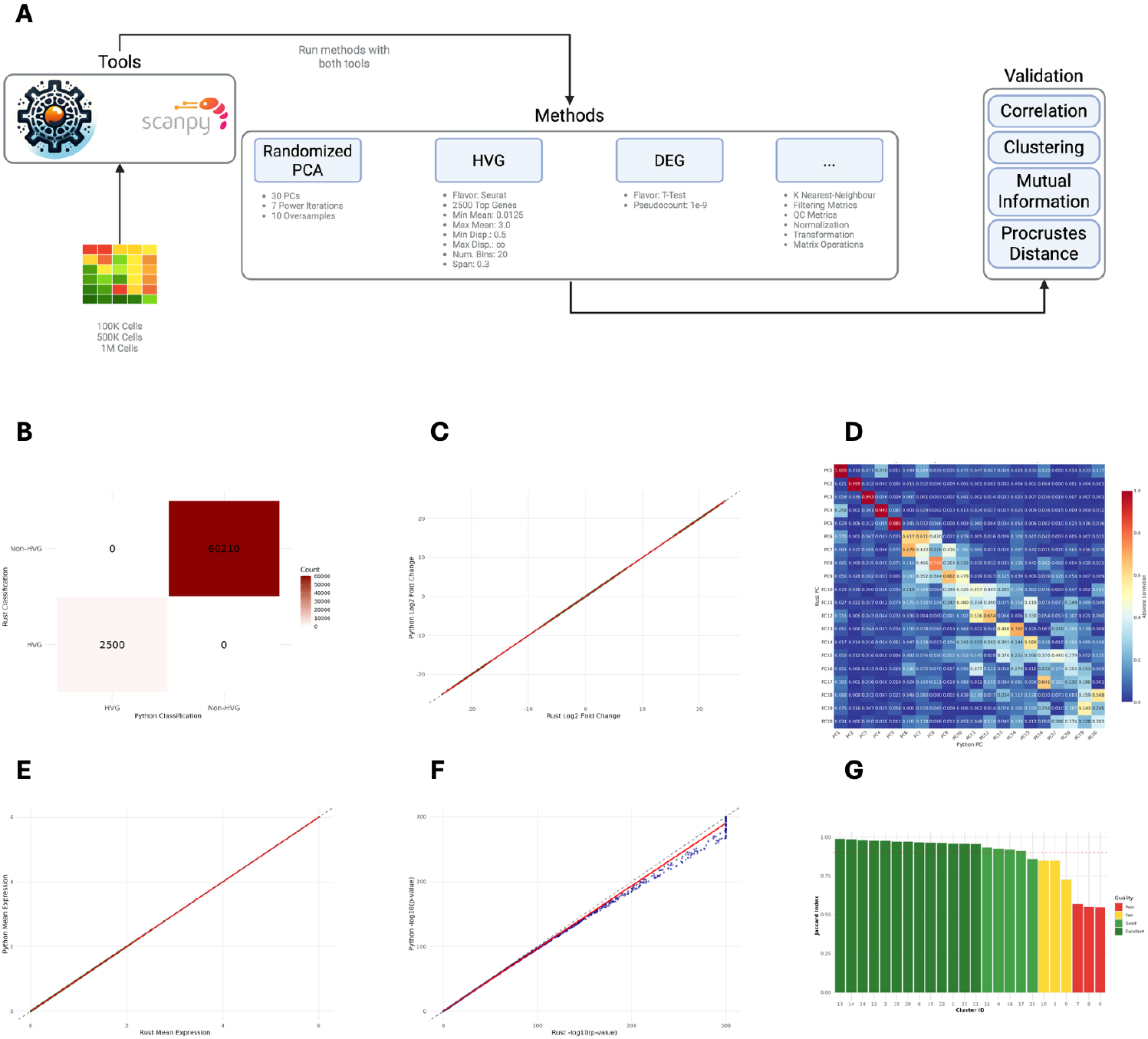
Validation of SingleRust methods against established implementations. **(A)** Validation workflow comparing SingleRust and Scanpy across three dataset sizes (100k, 500k, 1M cells) from the Tahoe dataset. Multiple computational methods were tested using correlation analysis, clustering concordance, mutual information, and Procrustes distance as validation metrics. **(B)** Overlap analysis of highly variable genes (HVG) showing perfect agreement (2,500/2,500 genes) between SingleRust and Scanpy implementations. **(C)** Scatter plot of log_2_ fold changes from differential expression analysis demonstrating near-perfect correlation (R^2^ = 1.0) between methods. **(D)** Correlation heatmap of principal components 1-20, showing strong agreement in the first 5 components (correlation ≥ 0.9) with expected decorrelation in higher dimensions due to randomized SVD initialization. **(E)** Perfect correlation of mean gene expression values across all tested genes. **(F)** P-value comparison from differential expression testing showing excellent agreement between implementations. **(G)** Clustering concordance analysis based on Leiden clustering of PCA embeddings, with high agreement (green bars) indicating consistent biological interpretation between methods.

**Fig. 4.**
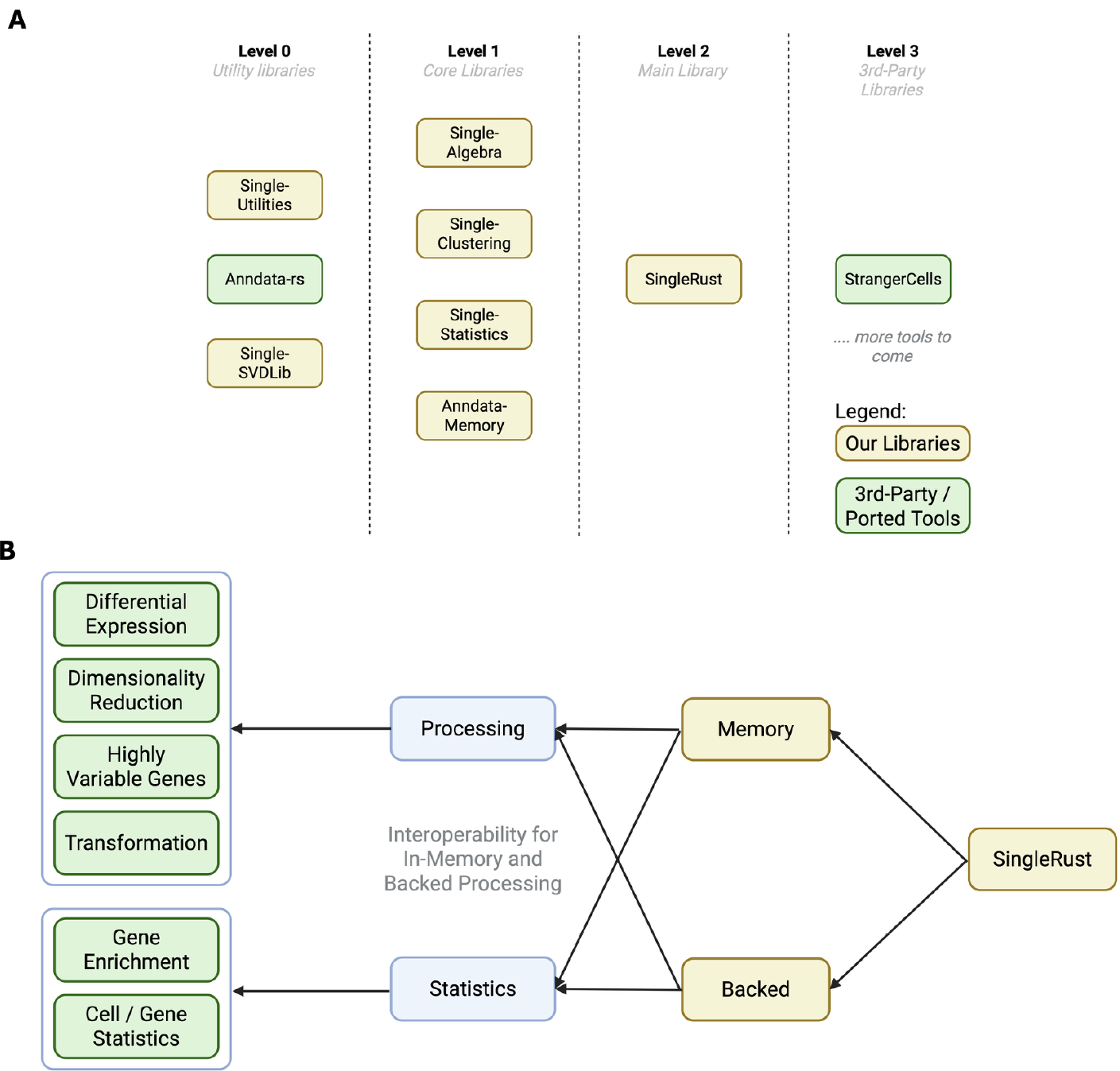
SingleRust library architecture. **(A)** Four-level modular hierarchy from utility libraries (Level 0) through core processing libraries (Level 1) to the main SingleRust library (Level 2) and third-party integrations (Level 3). **(B)** API structure with two high-level modules (Memory and Backed) sharing identical submodules for Processing and Statistics, ensuring consistent functionality across both execution modes.

### DEG Validation

Our 25.5x faster differential expression analysis maintains near-perfect agreement with Scanpy’s results. Log_2_ fold changes showed perfect correlation (*R*^2^ = 1.0, Figure 3C), while p-values demonstrated excellent agreement (Figure 3F). Minor deviations appeared only for genes with extremely low expression—likely reflecting differences in numerical precision handling between Rust and Python rather than algorithmic differences. These results confirm that SingleRust’s parallelized implementation produces statistically equivalent outputs while dramatically reducing computation time.

### PCA Validation

Despite implementing a memory-efficient masked sparse PCA algorithm, SingleRust maintained numerical concordance with Scanpy. The first five principal components showed correlations ≥ 0.90 (Figure 3D), while components 6-20 exhibited decreasing correlations (0.70-0.30), consistent with documented behavior of randomized SVD methods [10]. Functional validation through Leiden [11] clustering of PCA-reduced data demonstrated 96.0% agreement in cell assignments between implementations (n = 1,000,000 cells, Figure 3G). The 3.94% discordant assignments occurred predominantly at cluster boundaries, where minor variations in coordinate space can alter cluster membership. These results confirm that algorithmic optimizations for memory efficiency do not compromise biological interpretation.

These validation results demonstrate that SingleRust’s architectural advantages—zero-copy semantics, true parallelization, and optimized sparse operations, achieve substantial performance gains without sacrificing the numerical accuracy required for reliable biological discovery.

## 3 Discussion

### 3.1 Technical Architecture Enables Population-Scale Analysis

SingleRust demonstrates that 30 million cell analyses are feasible on standard 512 GB infrastructure, effectively doubling the practical capacity relative to current Python-based tools. This improvement results from systematic application of systems programming principles to single-cell algorithms.

The observed 2.4-25.5-fold performance improvements can be attributed to four architectural features. First, zero-copy semantics avoid the object duplication inherent in Python’s memory model [12], particularly beneficial for sparse matrix operations where redundant copies substantially increase memory pressure. Second, lock-free parallelization utilizes available CPU cores without Global Interpreter Lock constraints[13], enabling true concurrent execution. Third, compile-time type checking and memory safety [14] reduce runtime failures, though quantitative reliability improvements require longer-term evaluation. Fourth, deterministic execution facilitates reproducible analyses, addressing a key requirement for clinical translation [3].

These performance gains vary by operation type, reflecting underlying implementation architectures. For PCA, our 2.4-fold improvement represents Rust competing against optimized C++ code, as Scanpy delegates to pre-compiled NumPy/BLAS [15, 16] libraries with native multithreading, demonstrating that Rust’s memory model provides advantages even against mature compiled implementations. Conversely, operations constrained by Python’s Global Interpreter Lock show larger gains: differential expression analysis accelerates 24.6-fold through unrestricted parallelization across sparse matrix columns. The k-nearest neighbor implementation presents a particularly instructive case: despite Scanpy’s use of Numba-compiled kernels [8, 17], we observed severe cold-start penalties (25 seconds initial execution versus 2.2 seconds warm for 10,000 cells). This latency stems from Numba’s just-in-time compilation model, which must generate and optimize machine code on first invocation. While effective for repeated executions, this approach proves unsuitable for production environments requiring predictable performance from first call—containerized deployments, serverless functions, and batch processing pipelines all suffer from cumulative JIT overhead [18]. SingleRust’s ahead-of-time compilation eliminates this variability, providing consistent sub-second execution regardless of initialization state. Our results thus demonstrate that compilation strategy, not merely the presence of compiled code, determines production suitability for computational biology pipelines.

Memory efficiency improvements of 1.3-3.0-fold reflect complementary optimizations. The absence of garbage collection eliminates unpredictable memory overhead [19], while Rust’s ownership model enables in-place algorithms without defensive copying. These efficiencies scale with dataset size: our results show that operations requiring 450 GB for 10 million cells in Scanpy complete within 250 GB in SingleRust, enabling analysis of proportionally larger datasets on identical hardware. Notably, even at 30 million cells, SingleRust remained well within system capacity, HVG identification consumed 463 GB, PCA utilized 487 GB, and KNN peaked at 466 GB on our 512 GB system, maintaining 20-40 GB of available memory. This headroom contrasts sharply with Scanpy, which approached memory exhaustion at 10 million cells for several operations. The consistent 1.3-fold average memory reduction across all operations, while modest per operation, provides the cumulative margin necessary for population-scale analyses.

Despite achieving substantial memory reductions, our sparse matrix implementation remains constrained by current Rust ecosystem limitations. The nalgebra-sparse library enforce 64-bit integer indexing regardless of matrix dimensions, resulting in approximately 33% excess memory consumption compared to theoretical minimums. For typical single-cell matrices with fewer than 65,536 features, 16-bit indices would suffice, while many filtered datasets could utilize 8-bit indexing. We are developing an adaptive index representation that selects integer width based on matrix dimensions at runtime. This optimization could reduce index storage requirements by 4-8 fold, with particular benefit for the highly sparse matrices (¿95% zeros) characteristic of single-cell expression data. Initial implementations suggest this approach could support 30-35 million cell analyses on 512 GB systems, though we observe measurable degradation in random access patterns due to index decompression overhead. These trade-offs remain under active investigation. Importantly, SingleRust’s abstraction layers isolate such implementation details from the public API, enabling future deployment of optimized representations without disrupting existing workflows.

We are currently expanding our benchmarks to datasets exceeding 30 million cells to empirically determine SingleRust’s maximum capacity on standard infrastructure, with preliminary results suggesting viable processing up to 32.5 million cells for core operations. However, memory scaling remains fundamentally linear with cell count. Datasets exceeding 50 million cells will require architectural innovations beyond optimization—out-of-core processing, distributed computing, or novel data structures with sublinear memory complexity.

Our results establish that implementation-level improvements can extend the analytical frontier to 30 million cells, but approaching billion-cell datasets will demand algorithmic breakthroughs rather than incremental efficiency gains.

### 3.2 Numerical Validation Confirms Scientific Reliability

Computational efficiency without numerical fidelity would compromise scientific utility. We therefore conducted systematic comparisons between SingleRust and Scanpy implementations to assess whether our architectural optimizations, particularly shared-memory parallelism, maintain the analytical accuracy required for biological discovery.

Deterministic algorithms demonstrated complete concordance between implementations. The Seurat method for highly variable gene selection yielded identical results, with both tools selecting the same 2,500 genes from test datasets. Differential expression analysis produced equivalent statistical outputs, with log-fold changes showing perfect linear correlation (R^2^=1.0, n = 10,000 genes). P-values exhibited minor deviations only for genes with expression values below machine epsilon, where differences in floating-point arithmetic implementations between Rust and NumPy become apparent.

Stochastic algorithms exhibited variation consistent with theoretical expectations. Principal component analysis showed strong correlation for biologically relevant components, with components 1-5 achieving correlation coefficients ≥ 0.9. Higher components displayed expected decorrelation due to random initialization in truncated SVD algorithms, a well-characterized phenomenon in randomized linear algebra [10]. Importantly, this mathematical variation did not propagate to biological interpretation: Leiden clustering of PCA-transformed data assigned 96.06% of cells to identical clusters between implementations, with discordant assignments concentrated at cluster boundaries where stochastic effects are expected.

These results validate our core technical approach: leveraging Rust’s shared-memory model with read-write locks (RwLock) enables multiple threads to read sparse matrices concurrently without data duplication, while maintaining single-writer exclusivity for modifications. This zero-copy parallelism—impossible in Python due to the Global Interpreter Lock—allows us to accelerate computationally intensive operations without the memory overhead of thread-local copies. The validation data confirm that such architectural optimizations can coexist with numerical precision, suggesting that strategic use of compiled languages for specific computational bottlenecks represents a viable path for scaling single-cell analysis while maintaining compatibility with the broader Python ecosystem.

### 3.3 Practical Integration and Future Development

SingleRust facilitates adoption through systematic design decisions that prioritize compatibility with existing workflows. The library maintains full AnnData format compatibility [20] and implements APIs consistent with Scanpy’s [8] function signatures, enabling incremental adoption without disrupting established pipelines. This design enables researchers to selectively replace computationally intensive operations while retaining their existing analysis frameworks.

We provide two implementation strategies addressing distinct use cases. Researchers requiring fine-grained control can integrate individual SingleRust modules into existing Python workflows, leveraging performance improvements for specific bottlenecks. Alternatively, standardized pipeline implementations offer validated end-to-end workflows with predetermined parameters based on published best practices [21]. Both strategies benefit from static compilation, ensuring reproducible performance characteristics across computing environments and eliminating version-dependent behavioral variations common in interpreted languages.

Future development priorities reflect computational bottlenecks identified through community surveys and performance profiling. Planned Python bindings via PyO3 [22] will enable direct access to compiled algorithms from Python environments, with preliminary prototypes suggesting order-of-magnitude performance improvements for sparse matrix operations. This future interoperability will support a hybrid computational model where performance-critical algorithms execute in compiled code while maintaining Python’s advantages for interactive analysis and visualization integration.

Current implementation boundaries reflect deliberate prioritization of high-impact functionality. Graph-based visualization methods (UMAP[23], T-SNE[24, 25]) and trajectory inference algorithms remain unimplemented, while clustering performance requires additional optimization to match specialized implementations. This focused approach—addressing primary computational constraints before pursuing feature completeness—acknowledges that practical utility often derives from solving critical bottlenecks rather than comprehensive functionality. Community feedback through our public issue tracker informs quarterly development cycles, ensuring alignment with real-world computational challenges.

### 3.4 Implications for the Field

The transition of single-cell technologies from exploratory research to clinical diagnostics necessitates a fundamental shift in computational infrastructure requirements. Clinical applications demand deterministic execution, auditable processing pipelines, and guaranteed completion times [26], characteristics poorly served by interpreted languages with non-deterministic garbage collection and runtime type systems. SingleRust’s compile-time guarantees and predictable memory allocation patterns address these requirements, though extensive validation in clinical settings remains necessary.

Current trajectory analyses suggest single-cell datasets will reach 10^9^-10^10^ cells within five years, driven by population-scale studies [27], spatial transcriptomics, and longitudinal patient monitoring. This three-order-of-magnitude increase from current datasets will require fundamental algorithmic innovations beyond simple performance optimization. Our results demonstrate that systems programming languages can provide intermediate-term solutions, extending the useful lifetime of existing algorithms while the field develops genuinely scalable approaches such as geometric sketching or learned index structures.

The single-cell software ecosystem benefits from methodological diversity rather than monolithic solutions. Python-based tools excel at rapid prototyping and interactive visualization, while compiled languages like Rust address production deployment and performance-critical pipelines. This specialization enables researchers to select tools matched to specific computational requirements—leveraging Python’s flexibility for exploratory analysis and Rust’s efficiency for resource-intensive operations.

Our pure Rust implementation demonstrates the feasibility of compiled-language solutions for single-cell analysis, though current adoption requires researchers to work entirely within the Rust ecosystem. This requirement, while limiting immediate accessibility, enables us to fully explore the performance boundaries of systems programming approaches without the compromises inherent in cross-language interfaces. Future Python bindings will lower adoption barriers, but the current implementation serves as a crucial proof-of-concept that compiled languages can address the scale and reliability requirements of next-generation single-cell studies. Community feedback on this pure-Rust version will inform the design of future interoperability layers, ensuring that eventual Python integration preserves the performance characteristics demonstrated here.

## 4 Methods

### 4.1 Library Design and Implementation

We selected the Rust programming language for its unique combination of memory safety without garbage collection, zero-cost abstractions, and fearless concurrency model, which eliminate common runtime errors and enable predictable performance. SingleRust employs a modular ecosystem design with a three-tier architecture that separates concerns and enables independent development of components.

At the foundation (Level 0), single-utilities provides common traits and utility functions, while single-svdlib implements optimized singular value decomposition algorithms specifically tailored for sparse matrices. The data layer (Level 1) consists of anndata-rs (developed by Kai Zhang [28]) for backed data storage and reading / writing capabilities, complemented by our anndata-memory implementation that provides optimized in-memory representation with concurrent access via RWLocks. The processing layer (Level 1) includes single-algebra for matrix operations and PCA, single-statistics for statistical tests and multiple testing correction, and single-clustering for neighborhood search and clustering algorithms. At the user interface level (Level 2), the SingleRust library abstracts implementation details while providing a Scanpy-compatible API and fine-grained resource control options.

Thrid-party dependencies are carefully integrated to maximize performance: rayon provides work-stealing parallelization, parking lot offers efficient synchronization primitives, nalgebra and ndarray handle linear algebra operations, and simba enables SIMD vectorization. The type system design uses genrics for maximum flexibility and enum-based data storage following Rust best practices for heterogeneous data types.

Our memory management strategy prioritizes zero-copy semantics wherever possible, implements in-place type conversions for precision upgrades, and ensures explicit resource cleanup for predictable memory usage. The concurrent processing architecture features RWLock-based concurrent read access, a single-writer model for data safety, and rayon-based parallel iteration within read locks. We support dual processing modes: in-memory for performance-critical operations and backed/chunked processing for datasets exceeding available memory. The API design philosophy maintains Scanpy compatibility for user familiarity while exposing Rust-specific optimizations through optional parameters. Finally our extension and integration framework provides low-level access for third-party libraries wanting to integrate into the SingleRust ecosystem.

### 4.2 Core Algorithm Implementations

Our quality control metrics and filtering algorithms are fully multithreaded to maximize resource utilization. Additionally, our filtering implementation marks relevant cells and genes first, then executes a single subsetting operation. While this represents one larger operation moving more data, our performance evaluations demonstrate it outperforms the multiple separate calls approach used by existing tools.

For normalization, we followed the general implementation of Scanpy to calculate normalization factors for each cell, but extended the functionality to also allow gene normalization and implemented multi-threading for this operation. Additionally, we implemented log normalization for variance stabilization, providing users with more options for data processing.

Highly variable gene identification was implemented using Scanpy functions as a baseline while leveraging the optimized operations implemented in single-algebra. All three major flavors, Seurat[29], Cell Ranger[30], and SVR, are fully implemented, ensuring compatibility with different analytical approaches.

Dimensionality reduction has been implemented with SVD-based PCA. The most important improvement we implemented is sparse masked PCA to reduce memory consumption. Althouth this increases the possibility of cache misses, we observed a dramatic reduction in memory usage. We have implemented SVD using both the Lanczos algorithm and Randomized PCA [10, 31, 32] (similar to the scikit-learn implementation). For sparse-dense matrix multiplication operations, we developed our own custom algorithm capable of operating with masked sparse data structures. Additionally, in contrast to Scanpy, we implemented randomized PCA with mean centering to enable centered results production. The most important different from the Python version is how we calculate explained variance: while Python considers all features (all genes), we only consider the non-masked features during execution. We aim to provide users with options for computing both approaches in the future.

Differential expression analysis was implemented using our custom statistical testing library, optimized for testing on sparse data structures. Tests are executed in a multi-threaded manner, and we have implemented standard testing methods (T-Test and Mann-Whitney-U-Test) with multiple correction methods available. Additional testing methods are planned for future releases.

Our k-nearest neighbor (KNN) algorithm consists of two major components: KIDDO provides an exact KNN solution for search requests, while we have also implemented KNN using HNSW lib, which provides an efficient approximate KNN implementation in Rust based on the published algorithm [33]. We implemented an adaptive method that switches between KIDDO and HNSW based on dataset size. Our benchmarks show that at approximately 250, 000 cells, we observe a dramatic performance increase when switching to HNSW while maintaining similar accuracy to KIDDO. Both algorithms produce a CSR matrix with only the upper triangle stored, further reducing memory consumption. Currently, we have implemented only the Gaussian kernel method from Scanpy, as there is no UMAP Rust implementation yet available.

For clustering algorithms, we have implemented Leiden[11] and Louvain[34] as the primary methods. Both are still in alpha but already show improvements in terms of memory usage, though performance optimization remains ongoing. Both algorithms operate on a custom-implemented, multi-threading ready CSR network architecture, implemented in the single-clustering library.

Although we implemented these libraries with specific optimizations for single-cell data analysis, they are universally usable for other purposes as well, providing value beyond the single-cell community.

### 4.3 Benchmarking Methodology

All benchmarks were conducted on a medium-sized server equipped with a 64-core (128 threads) CPU, 512 GB of RAM, and an 8 TB SSD. We ensured a quiet system environment during all tests to minimize external inference with performance measurements.

For our test datasets, we selected the Tahoe-100M cells dataset [4] and created sub-sampled datasets of varying sizes (10K, 50k, 100k, 150k, 250k, 500k, 750k, 1M, 2.5M, 5M, 7.5M, 10M, 12.5M, 15M, 17.5M, 20M, 22.5M, 25M, 27.5M, and 30M cells). We applied random sub-sampling and merged different plates to produce the exact number of cells needed for each benchmark operation. These datasets were preprocessed, stored, and consistently used across all tests at their respective sizes.

We utilized Scanpy version 1.11.1 with Python 3.12.10 in a fresh environment to ensure reproducible comparisons. Parameters where matched as closely as possible between method calls in both Python and Rust implementations to ensure fair comparisons.

For CPU benchmarks, we executed each operation multiple times with three warmup runs followed by the actual measurement. When memory permitted multiple dataset copies, we conducted consecutive tests and measured execution time internally, which provided higher accuracy than external measurements. For memory-intensive mutating operations such as filtering or normalization that prevented multiple in-memory copies, we reloaded the data after each measurement to ensure consistent starting conditions.

All benchmarks were repeated multiple times to verify consistency between runs. When benchmarking complete pipelines, we executed the entire workflow several times (Python scripts for Scanpy, compiled binaries for SingleRust) but excluded the data loading step from measurements due to high variance between runs, likely caused by underlying system processes. Despite this variance, we observed no significant differences in loading performance between Scanpy and SingleRust.

Memory benchmarks followed the same methodology as pipeline benchmarks, with multiple executions of the complete binary to capture memory usage patterns. Statistical analysis of all benchmark results was performed using R to ensure rigorous evaluation of performance differences.

### 4.4 Numerical Validation Framework

We performed systematic validation of SingleRust implementations against scanpy using datasets of 100k, 500k, and 1M cells subsampled from the Tahoe-100M dataset. All datasets we stored in H5AD format to ensure consistent input data across both platforms.

#### Validated Operations

The following operations were validated across all dataset sizes:

- Highly Variable Gene (HVG) identification
- Principal Component Analysis (PCA)
- Differential Gene Expression (DEG)
- k-Nearest Neighbors (KNN)
- Normalization (library size and log1p)
- Cell and gene filtering
- Basic matrix operations (sum, mean, variance)

#### Validation Procedure

Each operation was executed in both Python (Scanpy) and Rust (SingleRust) environments with identical or equivalent parameters. Where exact parameter matching was not possible due to implementation differences, we selected the closest functional equivalents. All function calls used consistent random seeds where applicable.

##### Validation Metrics

Results were compared using the following metrics as appropriate for each operation:

- **Correlation**: Pearson and Spearman coefficients for continuous values
- **Clustering concordance**: Agreement of downstream clustering results
- **Mutual information**: Information-theoretic similarity measures
- **Procrustes distance**: Geometric alignment of embeddings
- **Adjusted Rand Index**: Clustering similarity accounting for chance
- **Jaccard Index**: Set overlap for discrete selections
- **Distance metrics**: Euclidean and Manhattan distances for vector comparisons

Tolerance thresholds were set based on operation type: exact agreement for discrete operations, relative tolerance of 10^−6^ for floating-point operations, and statistical similarity measures for stochastic algorithms.

## 5 Conclusion

SingleRust demonstrates that systems programming can address immediate scalability barriers in single-cell analysis while maintaining numerical fidelity. By reimplementing core algorithms in Rust, we achieved 2.4-25.5x performance improvements and enabled analysis of 30 million cells on standard infrastructure—effectively doubling current practical limits.

These gains, while substantial, represent an intermediate solution. As datasets approach billion-cell scales, algorithmic innovation must complement implementation optimization. SingleRust provides a high-performance foundation for exploring hybrid approaches: GPU acceleration for amenable operations, streaming algorithms for memory-constrained analyses, and novel data structures with improved complexity bounds.

The single-cell field stands at a computational crossroads. Clinical applications demand reliability and scale that current tools struggle to provide. By establishing that compiled languages can meet these requirements without sacrificing biological accuracy, this work opens new avenues for computational development. Future progress will likely emerge from combining the rapid innovation of interpreted languages with the performance guarantees of systems programming, a synthesis that SingleRust helps enable.

## Code Availability and Reproducibility

All code, including performance evaluation and numerical validation, is available at https://github.com/SingleRust and on our website https://singlerust.com.

## Acknowledgments

This study was financed through the DFG project 469073465, and the M.J. Fox Foundation (MJFF-021418; A.K. & T.W-C.). Figures were created with BioRender.com.

## Author contribution

I.F.D. Conceptualization, Implementation, Validation & Draft, M.F. Draft, Figure Design & Supervision, A.K. Draft revision & Supervision.

## Notes

https://github.com/singlerust

### Competing Interest Statement

The authors have declared no competing interest.

### Summary of Updates

in this revision, we expanded our benchmarks to 30 million cells, demonstrating that SingleRust can handle datasets up to 3x larger than what Scanpy can process before hitting memory limits. This showcases SingleRust's superior scalability for large-scale single-cell analysis. Additionally, we corrected minor issues in our benchmarking scripts that improve the accuracy of our performance evaluations.

